# Final amendment: Ambiguous specification of EGFR mutations compounded by nil or negligible fragmented gene counts and erroneous application of the Kappa statistic reiterates doubts on the veracity of the TEP-study

**DOI:** 10.1101/150904

**Authors:** Sandeep Chakraborty

**Affiliations:** R - 44/ 1, Celia Engineers, T. T. C Industrial Area, Rabale, Navi Mumbai, 400701, India

## Abstract

**Final amendment note:** This paper had raised two issues - the error-prone classification and mistaken application of the Kappa statistic. The classification critique still holds, and is being taken up with other criticisms at http://www.biorxiv.org/content/early/2017/07/02/146134. The Kappa statistic was an error on my part since I had failed to see another page in Table S1. Please consider this pre-print closed.

**Original abstract:** The use of RNA-seq from tumor-educated platelets (TEP) as a ‘liquid biopsy’ source [1] has been refuted recently (http://biorxiv.org/content/early/2017/06/05/146134, not peer-reviewed). The TEP-study also mentioned that mutant epidermal growth factor receptor (EGFR) was ‘accurately distinguished using surrogate TEP mRNA profiles’, which is contested here. It is shown that only 10 out of 24 (a smaller sample set, original study has 60) non-small cell lung carcinoma (NSCLC) samples here has any expression at all. Even there the number of reads (101 bp) are [1, 4, 1, 14, 9, 1, 2, 19, 21, 6], and do not even add up to one complete EGFR gene (about 6000 bp). EGFR mutations have been painstakingly collated in www.mycancergenome.org/content/disease/lung-cancer/egfr. In stark contrast, the TEP study has no specification of the EGFR mutant used. The TEP study found EGFR mutations in 17/21 (81%), and EGFR wild-type in 4/39 (10%) for NSCLC samples (Table S7, reflected in Fig 3, Panel E in percentages). A major flaw is the assumption that a non “EGFR wild-type” is a “EGFR mutant” since cases zero with EGFR reads (which are almost half of the samples) could be either. The application of the Kappa statistic to this data is erroneous for two reasons. First, the Kappa statistic does not handle “unknowns”, as is the case for samples with zero expression. Secondly, ‘interobserver variation can be measured in any situation in which two or more independent observers are evaluating the same thing’ [2]. The 90% (Fig 3, Panel E) is just the percentage of samples (35/39) that are not “EGFT WT” in one observation. It is not qualified to be in the Kappa matrix, where it translates to 35, leading to a Kappa=0.707, which implies “substantial agreement” [2]. The other observation (looking for EGFR mutation) is in a different set. To summarize, this work reiterates negligible expression of EGFR reads in NSCLC samples, and finds serious shortcomings in the statistical analysis of subsequent mutational analysis from these reads in the TEP-study.

## Introduction

Tumor tissue biopsy, the gold standard for cancer diagnostics, pose challenges that include access to the tumor, quantity and quality of tumoral material, lack of patient compliance, repeatability, and bias of sampling a specfic area of a single tumor [3]. This has resulted in a new medical and scientific paradigm defined by minimal invasiveness, high-efficiency, low-cost diagnostics [4], and, whenever possible, personalized treatment based on genetic and epigenetic composition [5]. The presence of fragmented DNA in the cell-free component of whole blood (cfDNA) [6], first reported in 1948 by Mandel and Metais, has been extensively researched for decades, with extremely promising results in certain niches [7]. Additionally, cfDNA derived from tumors (ctDNA) [8] have tremendous significance as a cancer diagnostic tool [9], and for monitoring responses to treatment [10]. However, detection of ctDNA, and differentiation with cfDNA, remains a challenge due the low amounts of ctDNA compared to cfDNA [11].

Recently, tumor-educated blood platelets (TEP) were proposed as an alternative source of tumor-related biological information [1, 12]. The hypothesis driving the potential diagnostic role of TEPs is based on the interaction between blood platelets and tumor cells, subsequently altering the RNA profile of platelets [13,14]. The study showed using RNA-seq data that tumor-educated platelets (TEP) can distinguish 228 patients with localized and metastasized tumors from 55 healthy individuals with 96% accuracy [1]. As validation, this study reported significant over-expression of MET genes in non-small cell lung carcinoma (NSCLC), and HER2/ERBB2 [15] genes in breast cancer, which are well-established biomarkers. The TEP-study also mentioned that mutant epidermal growth factor receptor (EGFR) was ‘accurately distinguished using surrogate TEP mRNA profiles’ [1]. EGFR, a trans-membrane glycoprotein, is responsible for triggering several signal transduction cascades implicated in more aggressive tumor phenotypes [16–18].

Previously, the TEP-study was refuted by an analysis of a subset of the samples (yet to be peer-reviewed) based on the absence of evidence of MET-overexpression in NSCLC samples [19]. Here, it is demonstrated that EGFR reads are equally negligible in the NSCLC samples. Next, it is noted that the kind of EGFR mutants are not explicitly specified (there are several EGFR mutations implicated in NSCLC - www.mycancergenome.org/content/disease/lung-cancer/egfr). Lastly, the Kappa statistic used is shown to be flawed for several reasons. In summary, this work reiterates the shortcomings of the TEP-study, which proposes the use platelets as a source for “liquid biopsy” [1].

## Results and discussion

### Null or negligible fragmented gene counts of EGFR

A smaller subset (24 out of 60) of NSCLC samples were used here. A large number of samples have zero counts (14 out of 24) (Table 1). Furthermore, the number of reads in other samples are negligible. These are 101 bp reads, and do not even add up to one complete gene (Fig 1). The differentiation of mutants from wild-type (WT) is a challenging task from such scanty reads.

**Table 1:**
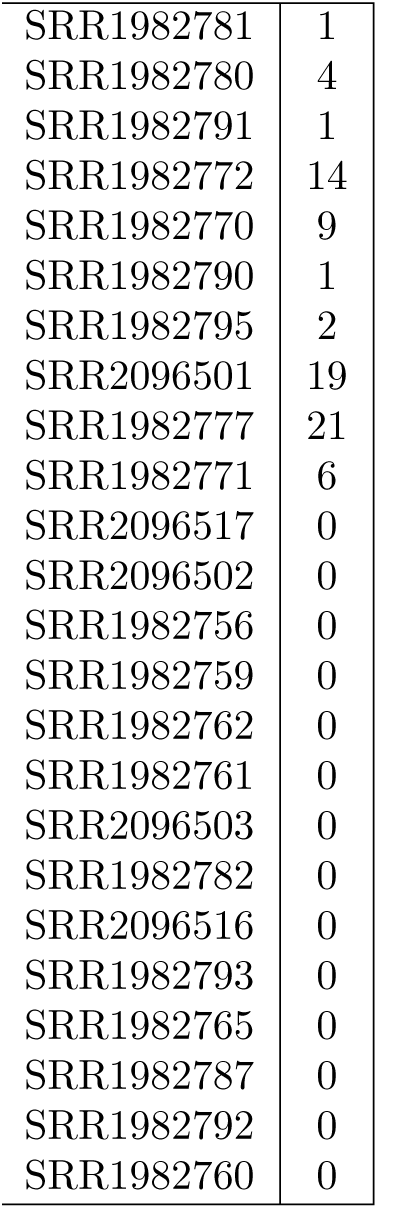
Counts of reads of the EGFR gene obtained using KEATS: A large number of samples have zero counts. Furthermore, the number of reads in other samples are negligible. Note, that these are 101 bp reads - so, they do not even add up to one complete gene. The differentiation of mutants from wild-type is a challenging task from such meagre counts.

|  |  |
| --- | --- |
| SRR1982781 | 1 |
| SRR1982780 | 4 |
| SRR1982791 | 1 |
| SRR1982772 | 14 |
| SRR1982770 | 9 |
| SRR1982790 | 1 |
| SRR1982795 | 2 |
| SRR2096501 | 19 |
| SRR1982777 | 21 |
| SRR1982771 | 6 |
| SRR2096517 | 0 |
| SRR2096502 | 0 |
| SRR1982756 | 0 |
| SRR1982759 | 0 |
| SRR1982762 | 0 |
| SRR1982761 | 0 |
| SRR2096503 | 0 |
| SRR1982782 | 0 |
| SRR2096516 | 0 |
| SRR1982793 | 0 |
| SRR1982765 | 0 |
| SRR1982787 | 0 |
| SRR1982792 | 0 |
| SRR1982760 | 0 |

**Table 2:** Kappa statistic derivation table for EGFR mutants and wild-type: The values are provided in Table S7 and also reflected in Fig 3, Panel E (which are percentages) in the TEP-study [1]. The computation of the Kappa statistic [2] corroborates a value of 0.707. However, the application of the Kappa statistic is erroneous since they involve mutually exclusive sets. Furthermore, in the presence of samples with absolutely no EGFR reads, this is not a binary situation wherein the absence of EGFR WT implies the presence of the EGFR mutant. The “35” (90% if you take percentages) is not where both observers agree that they are non-mutants (which gives the Kappa statistic such a good value of 0.707 which implies “Substantial agreement” [2]). It is just the percentage of one observer - it has no place in the Kappa matrix. One can make a matrix out of this data, but it can not be used to compute Kappa values.

|  | Second | Observer |  |
| --- | --- | --- | --- |
|  | Mut | Non-Mut | Total |
| First | 17 (81%) | 4 (10%) | 21 (m1) |
| Observer | 4 (19%) | 35(90%) | 39 (m2) |
| total | 21(n1) | 39(n2) | 60 (n) |

**Figure 1:**
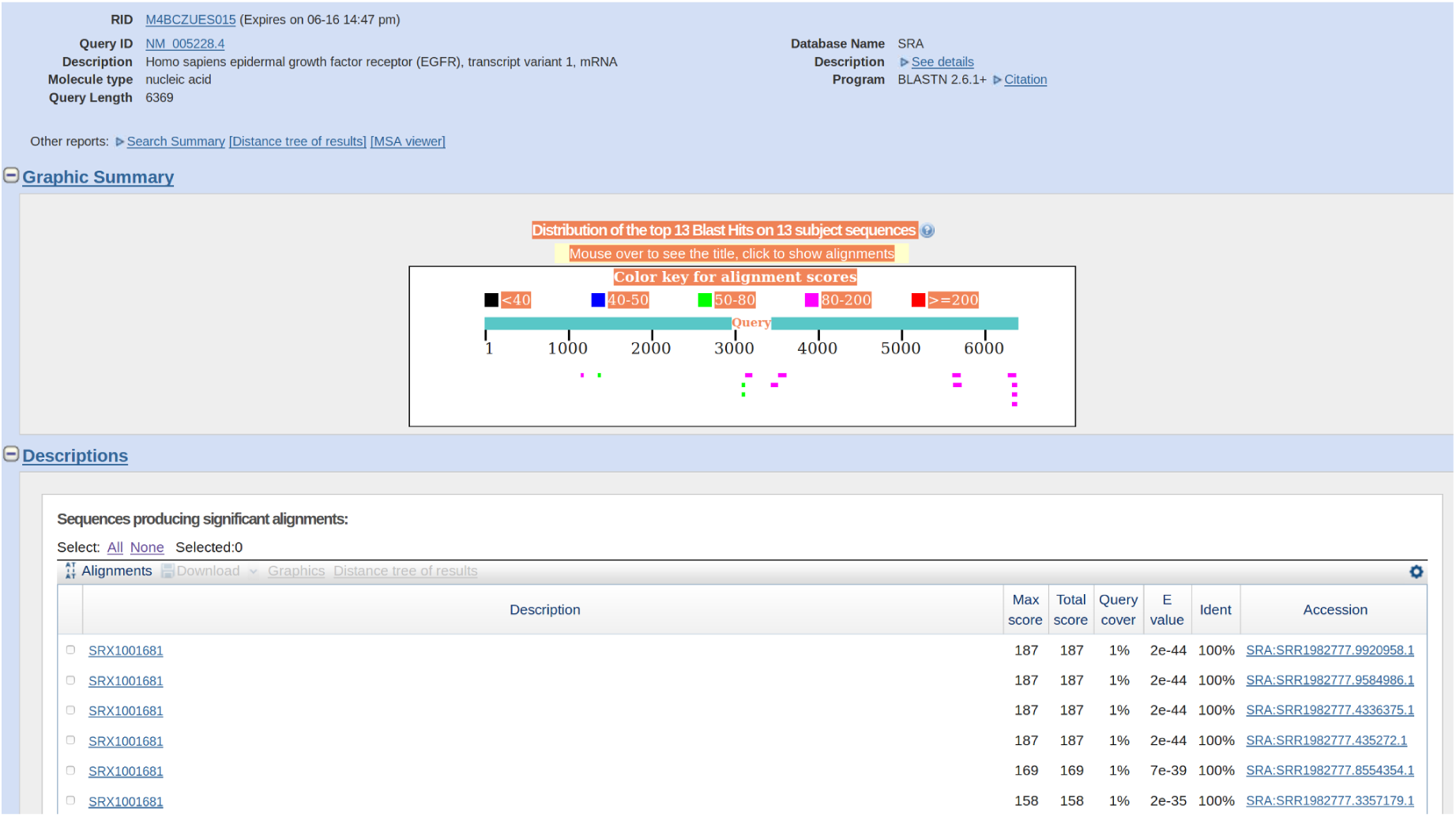
Low counts of EGFR reads in a NSCLC sample (SRA:SRR1982777) obtained using the online BLAST interface: NM_005228.4 (EGFR transcript variant 1, mRNA) was used as the query sequence. Identitying EGFR mutations from such a data is bound to be error-prone.

### Verification of the computation of the Kappa statistic for EGFR mutations

These formulas are obtained from [2].

pE = [(n1/n) * (m1/n)] + [(n2/n) * (m2/n)]

pE = (21*21 + 39*39) / 3600 = .545

pO = (17+35)/60 = 52/60 = 0.866

pO - pE = 0.866 - .545 = 0.32166

1 - pE = 1 - .545 = 0.455

Kappa = (pO - pE)/(1 - pE) = 0.32166 / 0.455 = 0.70694505494

Thus, this exactly corroborates the value (0.707) reported in Table S7 in the TEP-study [2].

### Erroneous application of the Kappa statistic in the EGFR study

However, the application of the Kappa statistic is seriously incorrect for two reasons.

1. The assumption that a sample that is not a “EGFR wild-type” is a “EGFR mutant” is seriously flawed in view of a large number of samples having zero EGFR reads (see above). The TEP-study found 4 “EGFR WT” in 39 samples, and automatically assumed that the rest 35 are EGFR mutants - this is clearly not correct for the samples with zero counts (Table 1). The Kappa statistic is a binary formula and does not handle a “do not know” scenario.
2. ‘Interobserver variation can be measured in any situation in which two or more independent observers are evaluating the same thing’ [2]. Here, the sets are apparently mutually exclusive with 21 and 39 elements, adding up to 60 NSCLC samples (although there is no explicit information).

To further emphasize the fuzziness, consider Fig 3, Panel E in [1] (which is essentially the Kappa matrix in percentages). The diagonal has 81% for both agreeing on “EGFR mutants” and 90% agreeing on “EGFR wild-type”. Agreement or disagreement does not makes sense if one is looking at different sets. The 90% is just the percentage of samples that are not “EGFT WT”. It is not agreement or disagreement - and thus, not qualified to be in the Kappa matrix.

## Conclusion

Statistical terminologies can often act as a smokescreen. Statistics needs to be supported by raw data. Just providing P-values and Kappa statistics can be misleading, and restrict future verification.

Absence of evidence is not evidence of absence in this case. In the absence of even a single reads, searching and failing to find a wild-type sequence does not imply the presence of a mutant. Furthermore, there is a flippancy in describing the EGFR mutant, since there can be many EGFR mutants - and the assumption that all NSCLC samples have the same mutation is biologically improbable.

This raises serious doubts on using TEP as a possible ‘liquid biopsy’ candidate. Essentially, it refutes the hypothesis that platelets carry enough RNA-seq from tumors to make it viable as a diagnostic method. A review found it ‘surprising’ that although ‘the tumor type was the predominant factor for the actual platelet conditioning, tumor metastasis did not significantly impact on them when compared to samples from patients without metastasis’ [14]. The excitement surrounding the fact that ‘2016 marked the first approval of a liquid biopsy test in oncology to assist in patient selection for treatment’ [20] should be tempered, and a cautious approach adopted [21, 22] with reports of ‘broken promises’ [23].

## Materials and methods

A kmer-based version (KEATS [24]) of YeATS [25–29] was used to obtain gene counts from transcripts in the RNA-seq data. A BLAST search suffices to demonstrate the absence of MET genes in the lung cancer samples. The Kappa statistic computation has been done based on formulas provided in [2].

## Competing interests

No competing interests were disclosed.

